# The effects of ploidy and mating system on the evolvability of populations: theoretical and empirical investigations

**DOI:** 10.1101/2025.07.17.665335

**Authors:** Carlos Olmedo-Castellanos, Diane Douet, Camilo Ferrón, Ana García-Muñoz, A. Jesús Muñoz-Pajares, Mohamed Abdelaziz, Josselin Clo

## Abstract

The amount of genetic diversity is a key parameter to understand the adaptive potential of populations. It has been demonstrated both theoretically and empirically that several factors influence genetic variance. In angiosperms, two of those are the ploidy level and the mating system of the populations. Polyploidy is theoretically known to increase adaptive potential in the long term. Self-fertilization has been theoretically associated with a decrease in genetic variance, even if it lacks empirical support. These factors have been studied independently, but are often shared in plants. However, there is a lack of empirical studies about the joint effects of polyploidy and selfing on genetic variance. In this paper, we conducted theoretical simulations to explore how genetic diversity could be affected by the ploidy level and mating system. We compared the simulation results with empirical estimates of genetic variance from the plant species *Erysimum incanum*, a selfing species from the Western Mediterranean basin exhibiting three different ploidy levels. We measured a series of phenotypic traits in individuals of each ploidy, obtained by controlled crosses and grown in different climatic conditions. While theoretical approaches showed a positive relationship between ploidy and genetic variance in both the short and long term, empirical results show lower evolvability and transgressive segregation for polyploids, both results being dependent on environmental conditions. Genetic variance in *E. incanum* polyploids could be related to recent establishment and adaptation to harsh environments, which explains the apparent contradiction with theory, where more settled and established populations are considered.

## Introduction

Polyploidy, the condition of having more than two sets of chromosomes, is widespread among plants and has long been recognized for its significant phenotypic effects, which are thought to play a key role in speciation and adaptation (Masterson, 1994; Soltis and Soltis, 1999; Otto and Whitton, 2000; Barker et al., 2016). Once viewed as an evolutionary dead end, recent studies suggest that polyploid species may be better equipped to cope with extreme and stressful environments (Bomblies, 2020; Van de Peer et al., 2021). However, polyploids, particularly young autopolyploid lineages, often exhibit higher extinction rates than their diploid counterparts (Mayrose et al., 2011; Arrigo and Barker, 2012; Levin, 2019), raising questions about their long-term adaptive potential. A possible explanation is that polyploid lineages could be associated with less genetic diversity, limiting their capacity to cope with environmental changes (Stebbins, 1966).

Nevertheless, theoretical studies suggest that polyploids may have greater heritable variance than their diploid relatives, especially over longer evolutionary timescales (Clo, 2022a). Several theoretical arguments support this view. First, due to higher levels of heterozygosity, recessive, weakly deleterious mutations are less purged and segregate at higher frequencies compared to diploid populations, thereby increasing genetic diversity (Otto and Whitton, 2000; Clo, 2022a). Second, non-additive genetic effects are expected to contribute more significantly to the heritable variance in polyploids than in diploids (Clo, 2022b). In particular, because two alleles per locus are transmitted through gametes, dominance variance (also called digenic interaction variance) becomes partially heritable and contributes to the adaptive potential of populations and species (Walsh, 2005). Finally, as polyploidy increases the potential for epistatic interactions, the influence of directional epistasis on genetic diversity is also predicted to be stronger (Mostafaee & Griswold, 2019; Clo, 2022b).

Empirical support for these theoretical predictions, however, remains limited. For example, Martin and Husband (2012) reported less heritable diversity in natural, established tetraploids compared to diploids in *Chamerion angustifolium*, while synthetic tetraploids had the highest amount of heritable genetic variance. This single study comparing genetic diversity for quantitative traits among cytotypes contradicts theoretical expectations. Other estimates of quantitative genetic variation in polyploids are scarce (O’Neil, 1997; Burgess et al., 2007), and generally fall within the range observed in diploid species (Clo et al., 2019). One possible reason for this mismatch is that theoretical models typically assume random mating, whereas polyploidy is frequently associated with shifts in mating systems, particularly toward increased self-fertilization (Barringer, 2007; Husband et al., 2008), which may influence the patterns of genetic variance.

Self-fertilization has a profound impact on genetic variation. By increasing homozygosity, any form of inbreeding increases genetic diversity (because you maintain the extreme homozygote genotypes and decrease the frequency of intermediate heterozygotes), but, over time, it also enables the purging of recessive deleterious mutations, thereby reducing genetic variance. Lande (1977), assuming a model of quantitative traits under stabilizing selection, found that selfing rates do not affect the genetic variance of traits under stabilizing selection, as increased homozygosity and purging counterbalance each other. Conversely, Charlesworth and Charlesworth (1995) concluded that high selfing rates strongly reduced genetic variance compared with fully outcrossing populations. More recent studies considering a wider range of selfing rates have reconciled these results, showing that the link between a decrease in genetic variance and the mating system is weak for most selfing rates, while for high selfing rates, a sharp decrease in genetic variance is observed (Abu Awad and Roze, 2018; Lande and Porcher, 2015). This reduction in genetic variance is highly sensitive to mutation rate, which leads to differences in the efficiency of purging, and is linked to genetic associations and the build-up and maintenance of linkage disequilibrium in selfing lineages. Indeed, these genetic associations contribute negatively to genetic variance, as it is a covariance between alleles of opposite effects (Abu Awad and Roze, 2018). These covariances can be potentially remobilized during rare outcrossing events, enabling the formation of new genetic combinations compared to parental genotypes (Clo et al., 2020). Empirical evidence indicates that the relationship between selfing and reduced genetic diversity is weaker than theoretical models suggest (Opedal et al., 2017; Clo et al., 2019). This difference between theoretical expectations and empirical results can be explained by the fact that theoretical models mainly consider an additive architecture, neglecting the effects of dominance on quantitative traits. Considering the effect of dominance, Clo and Opedal (2021) have shown that there is little difference in terms of heritable variance between selfing, mixed-mating, and outcrossing populations, which is more consistent with empirical findings. Notably, these insights are based solely on diploid systems, with no extension to polyploid contexts, to our knowledge.

The mainly Iberian–North African plant *Erysimum incanum* (Brassicaceae) offers a valuable system to investigate the joint effects of ploidy and mating system on genetic variance. This monocarpic, highly selfing species exhibits three ploidy levels (diploid, tetraploid, and hexaploidy) across its range in the Iberian Peninsula and Morocco (Nieto-Feliner, 1993; Abdelaziz et al., 2019; Abdelaziz et al., in prep.). It has a selfing mating system with a short and monocarpic life history, which facilitates working with several generations (García-Muñoz et al., 2023). Selfing, as well, is strongly promoted by *Anther rubbing* mechanism, which increases the self-pollination chances (Abdelaziz et al., 2019). Cytotypes are distributed across a latitudinal gradient, with diploids and tetraploids in the Iberian Peninsula and all three ploidy levels occurring in Morocco, from the Rif and Middle Atlas (diploids and tetraploids) to the Anti-Atlas (hexaploids), offering a natural context to examine how polyploidy and environment jointly shape genetic diversity.

In this study, we extended our theoretical and applied knowledge of the joint combination of mating system and ploidy level on the amount of heritable variance. To do so, we first generated theoretical expectations of how polyploidy and self-fertilization affect the amount and structure of genetic diversity of quantitative traits under stabilizing selection. Next, we estimated the amount of heritable variance of diploid, tetraploid, and hexaploid populations of *E. incanum*, in controlled and common garden conditions. Regarding theoretical modeling, we found that polyploidy does not qualitatively modify the expectation of the effect of the mating system on the amount and structure of the genetic variance. However, when looking at a fixed selfing rate, the higher the ploidy level, the higher the amount of genetic diversity compared to diploid populations. Empirical data showed, surprisingly, the opposite pattern, with diploids being generally the more genetically diverse cytotype, while tetraploids and hexaploids have lower genetic diversity.

## 2 Materials and Methods

### 2.1 Theoretical model

#### 2.1.1 General assumptions

A simulation model is developed to observe the evolution of a phenotypic trait *Z* in a population of size N_pop_. The model is an extension of Clo (2022a) in which we are now considering the mating systems of populations and higher ploidy levels (hexaploids). The selfing rate and the ploidy level are both parameters of the model. The considered populations can be diploids, tetraploids, or hexaploids, and each chromosome contains L loci. The selfing rate is a real number that can be set from 0 (obligate outcrossing) to 1 (fully selfing). The phenotypic trait value z of an individual is given by the following equation:

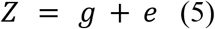

where e is a random environmental component drawn from a normal distribution with parameters 0 and V_e_, and g is the genotype of an individual computed as follows:

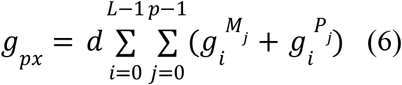

where *p* is the ploidy level, 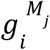 (resp. 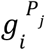 is the value stored in the locus *i* of the maternal (resp. paternal) chromosome *j*, and *d* is the dosage used to manage the effect of ploidy on genotypic values, set to 1 for diploid populations. In other words, the genotype of an individual is the sum of all values stored in its genome (full additivity at the phenotypic scale, but mutations are recessive at the fitness scale, see Clo, 2022a, for details), with a dosage factor for polyploid populations. We chose to focus on an additive model because it is much more commonly used than variation with non-additive effects (see Clo & Opedal, 2021, for a discussion), and then we have more points of comparison to judge how ploidy affects the outcome of the results. Also, adding non-additive effects when considering ploidy and inbreeding strongly complexifies the decomposition of the genetic variance, and up to twenty terms would be necessary for hexaploid populations (Gallais, 2003). The fitness of an individual with phenotype z is given by the following equation:

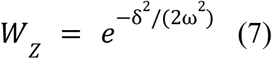

where δ is the difference between the phenotypic trait value z and an optimal trait value set to 0, and ω^2^ denotes the width of the fitness function, i.e., the strength of selection. All the notations and values used in the model and simulations are detailed in Table 1.

**Table 1.** Summary of notations and values.

| Symbol | Meaning | Values |
| --- | --- | --- |
| $N_{pop}$ | Population size | 250 |
| $L$ | Number of loci | 50 |
| $U$ | Mutation rate per haplotype | 0.1 or 0.005 |
| $a^2$ | Variance of mutational effects | 0.05 |
| $d$ | Dosage effect | 0.67 |
| $V_e$ | Variance of environmental effects | 1 |
| $\omega^2$ | Strength of selection | 1 or 9 |

#### 2.1.2 Individual-based simulations

At the beginning of the simulation, N_pop_ identical genomes are created with *p* chromosomes per individual, where *p* = 2, 4, or 6. The value initially stored in each locus is the optimum value 0. The simulation stops when an equilibrium for fitness is reached, more precisely when the quotient between two values of the mean fitness of the population, calculated 1000 generations apart, is greater than 99%.

The life cycle consists of three phases: selection, recombination, and mutation. A parent for the next generation is selected if the ratio between the fitness of the parent and the maximum fitness is greater than a randomly generated number between 0 and 1, drawn from a uniform distribution. If the selfing rate is higher than a new randomly generated number between 0 and 1, the same parent is selected again. Otherwise, a different parent is selected in the same way as described above. Each parent then passes on a gamete to the next generation, i.e. half of their chromosomes after recombination. Crossovers are modeled here as a random permutation between the alleles present at the same locus on homologous chromosomes. After gamete production, each haplotype can be subject to mutations. The number of mutations per haplotype is drawn from a Poisson distribution with parameter U, and their positions in the genome are randomly sampled from the L possible loci. The additive value of this mutation is then drawn from a normal distribution with parameters 0 and a^2^.

At the end of the simulation, the genetic variance V_g_, the genic variance V_a,_ and the genetic covariance COV_g_ at equilibrium are computed. The genetic variance corresponds to the variance between the complete genotypes, and the genic variance is the sum of the variances calculated at each locus separately. The three variables are linked by the following equation:

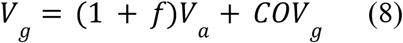

where *f* is the inbreeding coefficient of the studied population.

Inbreeding depression is also computed using the following formula:

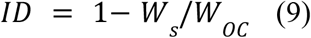

where W_s_ is the fitness of selfed individuals, and W_oc_ the fitness of outcrossed individuals in the population. These fitnesses are computed by generating 100 selfed and 100 outcrossed individuals from parents sampled at the mutation-selection-drift equilibrium.

To validate the results, the genetic variance obtained with the simulation considering a diploid and fully outcrossing population is compared with the stochastic house-of-cards approximation (formula 21.a) from Bürger et al. (1989). Using the notations from Table 1, this approximation can be rewritten as:

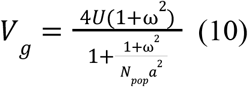

The value obtained with the simulation is close enough to the one obtained with the stochastic house-of-cards approximation to confirm the smooth running of the simulation.

#### 2.1.3 Choice of parameters

Simulations were run for different parameters. The haploid genomic mutation rate U is set to either 0.005 or 0.1, according to the range found in the literature (Keightley and Bataillon, 2000; Shaw et al., 2002; Halligan and Keightley, 2009). The parameters used for the number of loci L, the variance of mutational effects a^2^, the variance of environmental effects V_e_ and the strength of selection ω^2^ are similar to those used in Bürger et al. (1989); Ronce et al. (2009), and are the following: L = 50, a^2^ = 0.05, V_e_ = 1 and ω^2^ = 1 or 9 (see also Clo and Opedal (2021)). The dosage effect d is set to 0.67 (Porturas et al., 2019; Clo and Kolář, 2021). In each simulation, the population contains N_pop_ = 250 individuals. The parameter values are summarized in Table 1, but they have been selected to fit with simulations performed in Clo (2022a) and Clo *et al*. (2020).

### 2.2 Empirical investigation

#### 2.2.1 Study system

*Erysimum incanum* is a species of the Brassicaceae family from the Iberian Peninsula and Morocco, part of the INCANUM species complex. We can find populations of three ploidy levels: diploid, tetraploid, and hexaploid (n = 8 chromosomes; Nieto-Feliner, 1993; Abdelaziz et al., in prep.). This species complex is predominantly autogamous. Flowers are small, hermaphrodite, with tetradynamous stamens (four long and two short) typical in Brassicaceae, and self-compatible. Flowers also show the so-called anther rubbing mechanism (Abdelaziz et al., 2019), which strongly promotes selfing. Diploids show a vicariant distribution in the Rif and the Pyrenees Mountains, and tetraploids are localized in the south-western Iberian Peninsula and the Middle Atlas Mountains (Nieto-Feliner, 1993; Fennane et al., 1999). Hexaploid populations are situated in the southernmost ranges of Morocco (High Atlas and Anti-Atlas; Abdelaziz et al., in prep.).

#### 2.2.2 Experimental design and phenotypic measurements

Selfing and outcrossing populations were needed to properly infer the heritable variance and the capacity to generate new phenotypic combinations after recombination. To obtain selfing and outcrossing populations, we sowed seeds from wild populations of the three ploidies of *Erysimum incanum* in the common garden at the Faculty of Science of the University of Granada. We chose two populations within each ploidy to collect the seeds. We performed three generations of selfing to standardize maternal effects before carrying out the controlled crosses according to the experimental design. We then produced two types of controlled crossings: selfing (self-fertilized plants) and outcrossing (crossing two inbred lines within populations). Individuals were tagged according to each crossing, obtaining full-sibling groups (families). We finally obtained 155 diploid individuals (135 selfing in 30 families and 25 outcrossing in 4 families), 189 tetraploids (161 selfing in 52 families and 28 outcrossing in 6 families), and 104 hexaploids (66 selfing in 16 families and 38 outcrossing in 8 families). Groups were divided and separately exposed to three environments: greenhouse, common garden in the Faculty of Science of Granada (Spain), and mountain conditions in Hoya de Pedraza Botanic Garden in Sierra Nevada (Granada, Spain), to see the effect of the growing site on the measurement of genetic diversity. We did not expose outcrossing individuals to mountain conditions due to logistical problems during the experiment. Sample sizes are shown in Table S1.

We measured a series of morphological and floral traits. Morphological traits that we measured were the height of the main stalk, the number and diameter of all stalks, and the total number of flowers. Floral traits were petal length (maximal length from the edge of the petal to the curving point), corolla diameter (distance between edges of two opposite petals), and corolla tube length (distance from sepals’ basis to corolla aperture), the length of a long stamen filament and a short filament and the style height (from the basis of the style to the stigma), so that we could calculate herkogamy (i.e. vertical distance difference between long stamen and style). We took measurements in one generation of each controlled cross.

To obtain fitness information, we registered the number of fruits per plant, and we estimated the average number of seeds in each plant by counting seeds from four fruits. We then obtained the fitness value as the product of the total production of fruits and the total production of seeds per individual. In each ploidy, we used the highest value to standardize fitness data.

#### 2.2.3 Evolvability for selfing individuals

To measure the evolutionary potential of different ploidy groups, we calculated the environmental and genetic variances for the different phenotypic traits of the selfing groups at every ploidy level (diploid, tetraploid, and hexaploid; Table 2; see Table S2 for detailed results). Genetic variance was considered as the between-selfing group variance, while environmental variance was represented by the within-group variance. Total genetic variance is expected to be a better proxy of the evolutionary potential than additive variance for predominantly selfing species for several reasons (Sztepanacz et al., 2023). First, because predominantly selfing are made of several purely inbred lines that rarely intercross (Siol et al., 2008; Jullien et al., 2019; Clo et al. 2021), such that the short-term response to selection is made at the level of the inbred line (Lande & Porcher, 2015; Abu Awad & Roze, 2018; Clo & Opedal, 2021). Second, because some forms of dominance variance become heritable with inbreeding and polyploidy (Kelly, 1999; Walsch, 2005). It has been shown theoretically (Clo and Opedal, 2021) and empirically (Sztepanacz et al., 2023) that these new components can contribute substantially to the genetic diversity of quantitative traits. All these components are included in the among-lines variations, allowing an easy way to estimate the short-term heritable variance and to compare heritable diversity among cytotypes. It is only possible because *E. incanum* is a primary selfing species. Comparing heritable diversity among outcrossing populations of different ploidy levels is a more challenging task (see Clo, 2022b for a discussion).

**Table 2.**
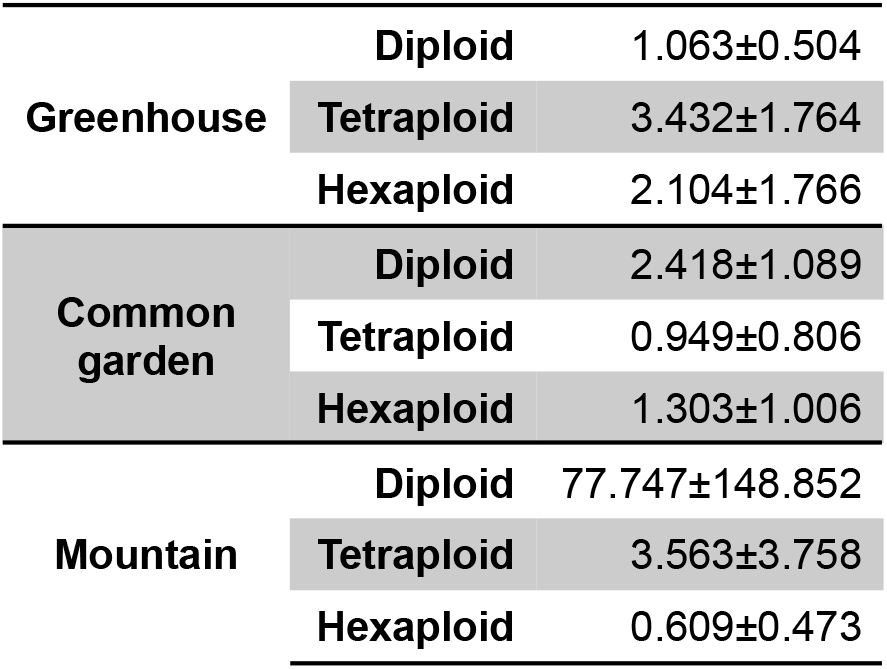
Evolvability means and standard deviations for each ploidy and environmental levels in individual and flower traits of *Erysimum incanum*.

We calculated evolvability as the ratio of the genetic variance VG for a given trait and its squared mean (*x*^2^) using the following formula from Houle (1992):

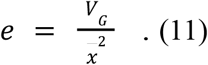

We also calculated the broad-sense heritability for each trait, as the ratio of the genetic variance over the phenotypic variance. Evolvability is a more accurate way to approach the adaptive potential of populations because the additive variance of a trait is correlated with both the environmental and epistatic variances (Carter et al., 2005; Hansen et al., 2011), such that the heritability of a trait can be close to zero while the trait exhibits substantial additive variance. We checked if ploidy had any effect on evolvability and heritability with Kruskal-Wallis tests as well as a t-test for both measurements. To extract genetic variance, we performed a general linear model test on the data to assess family-level effects using R 4.1.3 (Table 3).

**Table 3.** Statistics of general linear model of evolvability on ploidy and environment effects on individual and flower traits of *Erysimum incanum*. Greenhouse and diploid levels are used as default conditions. CG: Common garden; MT: Mountain.

|  | Estimate | SD | Error | t value |
| --- | --- | --- | --- | --- |
| **(Intercept)** | 1.063 | 16.548 | 0.064 | 0.94896 |
| **Tetraploid** | 2.369 | 23.402 | 0.101 | 0.91961 |
| **Hexaploid** | 1.042 | 23.402 | 0.045 | 0.96460 |
| **CG** | 1.355 | 23.402 | 0.058 | 0.95396 |
| **MT** | 76.684 | 23.402 | 3.277 | 0.00155** |
| **Tetraploid:CG** | -3.838 | 33.096 | -0.116 | 0.90796 |
| **Hexaploid:CG** | -2.157 | 33.096 | -0.065 | 0.94820 |
| **Tetraploid:MT** | -76.553 | 33.096 | -2.313 | 0.02326* |
| **Hexaploid:MT** | -78.180 | 33.096 | -2.362 | 0.02057* |

#### 2.2.4 Transgressive segregation

As mentioned in the introduction, recombination events among inbred lines can lead to the formation of new allelic combinations, potentially allowing for better response to selection in the long term. To test for this effect, we looked at transgressive segregation patterns (presence of outlier phenotypes in the recombining outcrossing populations compared to their selfing progenitors). We first looked at the minimum and maximum phenotypic values within the selfing and outcrossing groups for each ploidy level to apply the following classical transgressive segregation formula from Rieseberg et al. (2003, 1999):

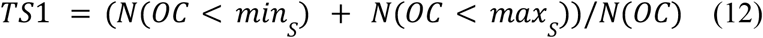

Where TS=Transgressive segregation; N=number, OC=outcrossing, S=selfing, max=maximum value for a trait measurement; min=minimum number for a trait measurement. We also assessed transgressive segregation using the index proposed by Koibe et al. (2019), by calculating the proportion of outcrossing individuals out of the range defined by the phenotype distribution on the selfing group:

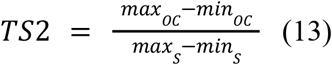

We finally calculated segregation variance of outcrossing offspring for each ploidy compared with the selfing group as the proportion of phenotypic variance from outcrossing offspring (V^P^ _os_) compared to selfing ones (V^P^ _s_) as follows:

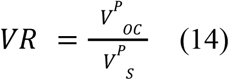

These ratios gave information about the novel phenotypes available in a group regarding the previous one, and the capacity of a group to generate these new phenotypes. This is an indirect way of evaluating mid-term evolutionary potential. We also performed a Kruskal-Wallis test to explore ploidy effects on transgressive segregation.

## 3 Results

### 3.1 Theoretical expectations

Figure 1 and Figures S1-S3 present trends in genetic variance, the genic variance, and the genetic covariance as functions of the selfing rate, under varying combinations of mutation rate (U) and strength of selection (ω^2^), as defined in Table 1. Across all parameter sets, both genetic variance and the genetic covariance decrease with increasing selfing rate, regardless of the parameters (Figures 1, S1-S3). Polyploid populations consistently exhibited higher genetic variance, higher genic variance, and higher genetic covariance in absolute value (as the covariance is mostly negative; Figure 1). The ratios between the genetic, the genic variance, and the covariance between polyploid and diploid populations tend to decrease overall as the selfing rate increases (Figures 1, S1-S3).

**Figure 1.**
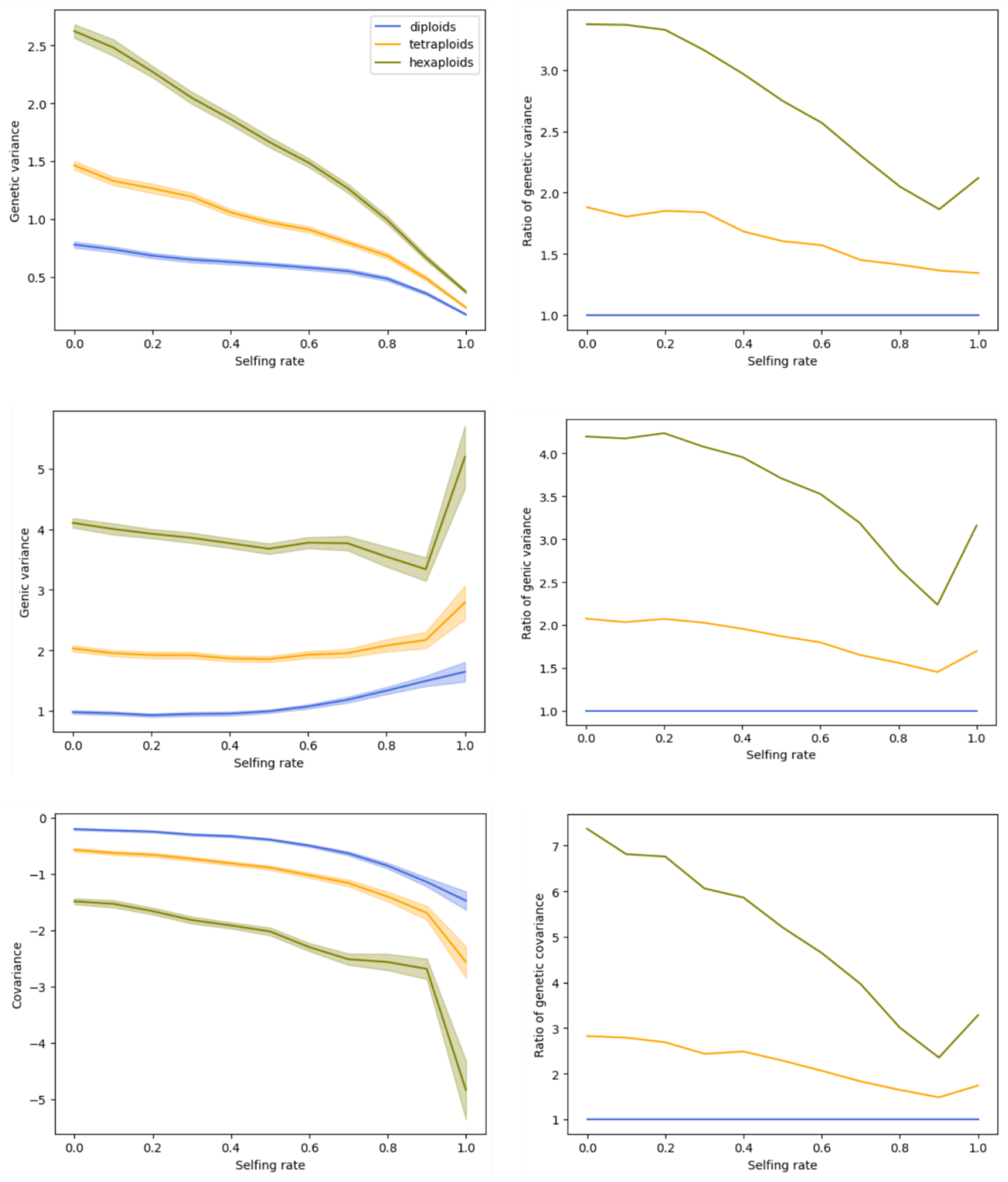
Genetic variance, genic variance and covariance and their 95% confidence intervals (left) and their ratios between polyploid and diploid populations (right) as functions of the selfing rate for different levels of ploidy, when *U* = 0.1 and *ω*^2^ = 1.

On the other hand, the variations of the genic variance as a function of the selfing rate depend on the haploid mutation rate U. Figures 1 and S2 indicate that for a mutation rate U of 0.1, the genic variance increases with the selfing rate. Figures S1 and S3 show that the genic variance is a decreasing function of the selfing rate when U = 0.005. Moreover, a higher mutation rate U leads to higher values of the genetic variance, the genic variance, and the genetic covariance in absolute value, for all ploidy levels and all selfing rates. Finally, the stronger the selection (smaller ω^2^), the lower the amount of genetic diversity in populations (see, for example, Figure 1 compared to Figure S2).

Regarding inbreeding depression, results from Figure 2 indicate a general decline in inbreeding depression with increasing selfing rate across all parameter combinations. Moreover, inbreeding depression was consistently more pronounced in polyploid populations compared to diploids (Figure 2).

**Figure 2.**
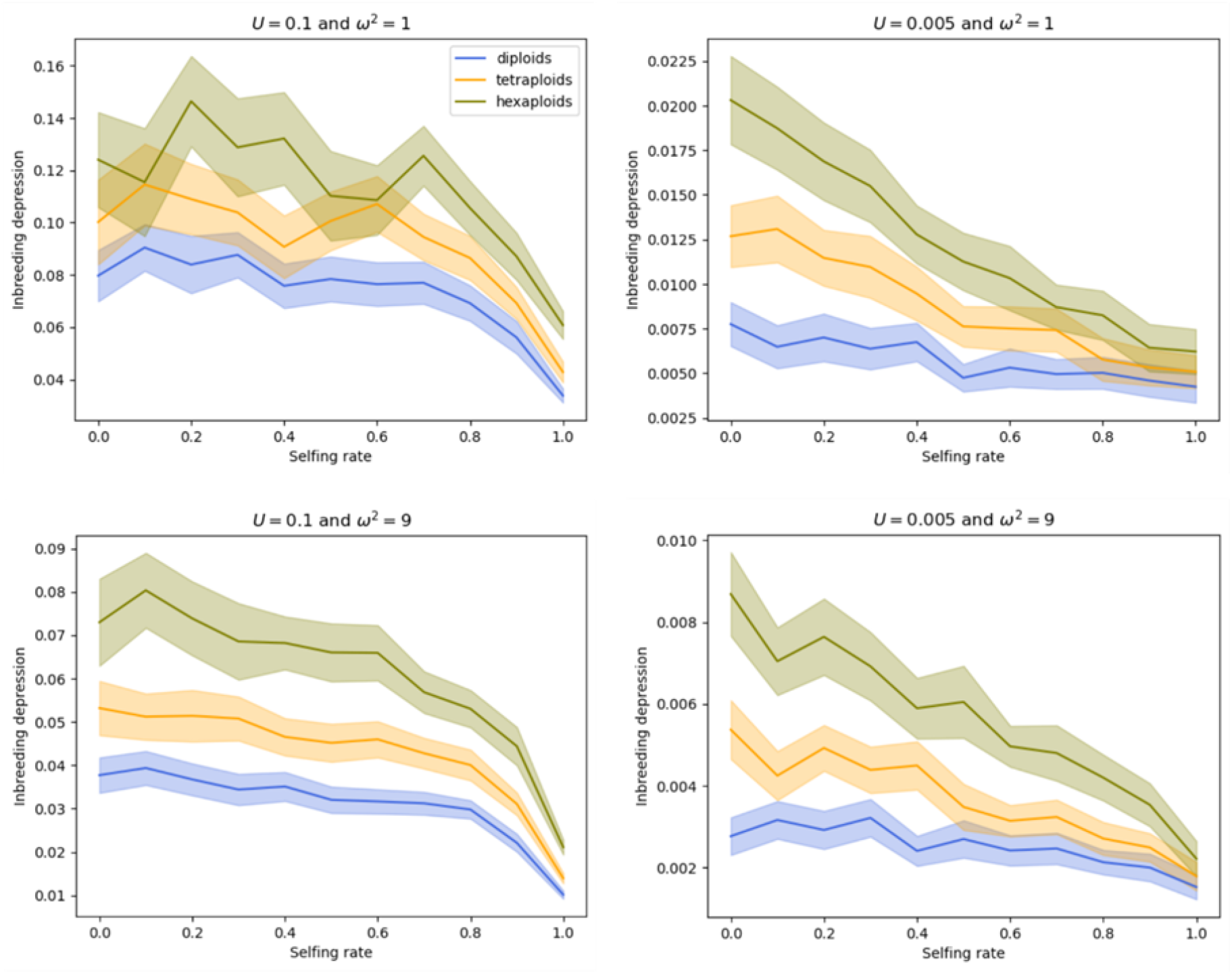
Inbreeding depression and associated 95% confidence intervals as a function of the selfing rate for different values of the parameters *U* and *ω*^2^, and for different levels of ploidy.

As an intermediate summary, we found theoretically that polyploid populations should have more genetic diversity than their diploid counterparts. We also found that polyploid populations stored more genetic diversity through genetic covariation and linkage disequilibrium, such that we should find transgressive segregation patterns more frequently in polyploids than in diploids.

### 3.2 Empirical data

#### 3.2.1 Evolvability for selfing individuals

We estimated evolvability for each phenotypic trait to assess the evolutionary potential across ploidy levels and environments. In both common garden and mountain settings, diploid populations consistently exhibited the highest evolvability levels (Tables 2 & 3). In contrast, tetraploid and hexaploid individuals showed significantly reduced evolvability under mountain conditions, but not under common garden conditions (Tables 2 & 3). Across all the ploidy levels, mountain conditions tended to increase evolvability (Tables 2 & 3). However, ploidy itself did not exert a statistically significant main effect on evolvability when considered independently (Tables 2 & 3).

#### 3.2.2 Transgressive segregation

We assessed transgressive segregation using three complementary indices (Table 4 for greenhouse and Table 5 for common garden): the classical index (TS1) by Rieseberg et al. (2003), the range-based index (TS2) from Koibe et al. (2019), and the variance ratio (VR). TS2 index and the variance ratio proved more sensitive to detecting transgressive phenotypes, whereas the classical Rieseberg index was more conservative.

**Table 4.** Transgressive segregation indexes for individual and phenotypic traits in *Erysimum incanum* in the greenhouse. We considered TS1 as the classical index proposed by Rieseberg et al. (1999, 2003) TS2 as the proposed index by Koibe et al., (2019) and variance ratio between outcrossing and selfing groups.

|  | Diploid |  |  | Tetraploid |  |  | Hexaploid |  |  |
| --- | --- | --- | --- | --- | --- | --- | --- | --- | --- |
|  | TS2 | TS1 | Variance ratio | TS2 | TS1 | Variance ratio | TS2 | TS1 | Variance ratio |
| **Plant height** | 0 | 0,512 | 0,584 | 0,021 | 1,234 | 1,034 | 0,016 | 0,922 | 0,9 |
| **Number of stalks** | 0 | 0,947 | 1,894 | 0 | 0,727 | 0,777 | 0 | 0,769 | 0,656 |
| **Stalk diameter** | 0,02 | 0,943 | 1,196 | 0,021 | 0,933 | 1,369 | 0,016 | 1,049 | 1,031 |
| **Number of flowers** | 0,061 | 0,538 | 0,93 | 0 | 0,713 | 0,748 | 0 | 0,947 | 0,791 |
| **Petal length** | 0,02 | 1,426 | 1,383 | 0,042 | 1,107 | 0,946 | 0 | 0,564 | 0,934 |
| **Flower diameter** | 0,061 | 0,927 | 0,782 | 0,021 | 0,973 | 0,841 | 0 | 0,882 | 0,998 |
| **Corolla tube length** | 0,041 | 1,013 | 1,454 | 0,083 | 1,06 | 1,503 | 0 | 0,931 | 1,286 |
| **Long filament length** | 0,082 | 1,168 | 1,338 | 0,021 | 0,727 | 0,855 | 0 | 0,872 | 0,875 |
| **Short filament length** | 0,041 | 1,093 | 1,149 | 0,021 | 0,85 | 0,879 | 0 | 0,699 | 0,799 |
| **Style length** | 0,02 | 0,854 | 1,048 | 0,021 | 0,801 | 0,879 | 0,016 | 0,909 | 1,341 |
| **Herkogamy** | 0,02 | 0,939 | 0,799 | 0 | 0,83 | 1,177 | 0,048 | 0,954 | 1,42 |

**Table 5.** Transgressive segregation indexes for individual and phenotypic traits in *Erysimum incanum* in the common garden. We considered TS1 as the classical index proposed by Rieseberg et al. (1999, 2003) TS2 as the proposed index by Koibe et al., (2019) and variance ratio between outcrossing and selfing groups.

|  | Diploid |  |  | Tetraploid |  |  | Hexaploid |  |  |
| --- | --- | --- | --- | --- | --- | --- | --- | --- | --- |
|  | TS1 | TS2 | Variance ratio | TS1 | TS2 | Variance ratio | TS1 | TS2 | Variance ratio |
| Plant height | 0 | 0,012 | 0 | 0,048 | 1,088 | 1,14 | 0,041 | 1,018 | 1,346 |
| Number of stalks | 0 | 0,333 | 0,201 | 0,048 | 2,167 | 4,999 | 0,02 | 1,667 | 1,849 |
| Stalk diameter | 0 | 0,674 | 0,541 | 0 | 0,598 | 0,715 | 0,02 | 1,296 | 1,177 |
| Number of flowers | 0,25 | 0,209 | 0,057 | 0 | 0,574 | 0,941 | 0,082 | 1,521 | 1,402 |
| Petal length | 0 | 0,204 | 0,105 | 0 | 0,466 | 0,543 | 0 | 0,621 | 0,801 |
| Flower diameter | 0 | 0,461 | 0,504 | 0,048 | 0,914 | 1,52 | 0 | 0,912 | 1,017 |
| Corolla tube length | 0 | 0,236 | 0,125 | 0 | 0,556 | 0,557 | 0,061 | 1,003 | 0,714 |
| Long filament length | 0 | 0,222 | 0,175 | 0 | 0,494 | 0,619 | 0 | 0,65 | 0,613 |
| Short filament length | 0 | 0,408 | 0,523 | 0 | 0,831 | 1,022 | 0 | 0,726 | 0,969 |
| Style length | 0 | 0,132 | 0,083 | 0 | 0,4 | 0,738 | 0 | 0,684 | 0,64 |
| Herkogamy | 0 | 0,077 | 0,023 | 0 | 0,663 | 1,112 | 0 | 0,637 | 0,838 |

General linear model analyses revealed that polyploidy level had a negative effect on transgressive segregation, while mountain conditions had a positive effect, particularly for the Koibe index and the variance ratio. In contrast, the Rieseberg index (TS1) showed no significant effects of ploidy, environment, or their interaction.

## 4. Discussion

In this study, we found that polyploidy is associated with reduced genetic diversity and a lower potential for transgressive segregation. These findings stand in contrast with our theoretical predictions, which suggest that, all else being equal, polyploidy should enhance heritable variance and increase the likelihood of generating transgressive phenotypes due to greater genetic covariance following recombination.

### Theoretical evolution of genetic diversity after polyploidization

Our simulations showed that the evolution in genetic diversity in polyploids qualitatively mirrors that in diploids. For all ploidy levels, both genetic variance and (mostly negative) genetic covariance declined with increasing selfing rates. Inbreeding depression also decreased with increasing selfing rates. However, at any given selfing rate, polyploids maintained higher genetic (co-)variance than diploids. These findings are consistent with theoretical expectations (Otto & Whitton, 2000; Clo, 2022a).

Notably, polyploids under full selfing exhibited lower genetic variance than diploids under full outcrossing, emphasizing the importance of mating strategies in shaping adaptive potential. While selfing consistently reduced genetic variance, this decline was gradual in diploids and became more pronounced only at high selfing rates, particularly when mutation rates were elevated—again aligning with prior studies (Abu Awad and Roze, 2018; Lande and Porcher, 2015). In this work, these results have been extended to polyploid populations. For polyploids, simulations showed that the decrease in genetic variance and covariance as functions of the selfing rate is faster than for diploids.

Interestingly, our simulations also highlight that inbreeding depression increased with the ploidy level, but decreased with higher selfing rates, regardless of ploidy. The latter reflects the increased homozygosity in selfing individuals, which facilitates purging of deleterious alleles (Ronfort, 1999). However, the greater inbreeding depression observed in polyploids compared to diploids is in contradiction with previous models (Lande & Schemske, 1985; Ronfort, 1999). This is likely due to the use of a quantitative genetics model that allows for genetic covariation among deleterious alleles that are more pronounced in polyploid populations, limiting the purge of those alleles. Our finding also contradicts some empirical studies suggesting that polyploids should exhibit equal or lower inbreeding depression than diploids (Clo & Kolář, 2022).

### How polyploidy affects evolvability in practice

Our empirical results revealed that evolvability differs across ploidy levels and environments (Table 2). Diploid populations consistently exhibited the highest evolvability under both common garden and mountain conditions. This aligns with previous work showing higher evolvability in diploids compared to naturally established tetraploids (Martin & Husband, 2012). On the contrary, other studies report similar or lower heritable diversity in polyploids (O’Neil, 1997; Burgess et al., 2007; Ebadi et al., 2023), partly supporting our findings and challenging broad generalizations derived from theory (Clo, 2022a).

Several factors may explain this discrepancy. Newly formed polyploids may experience reduced genetic diversity due to allelic dilution and post-polyploidization bottlenecks (Clo, 2022b; Mortier et al., 2024). Additionally, while selfing facilitates establishment in novel environments, it does not enhance genetic diversity as outcrossing does (Wright et al., 2008). Therefore, discrepancies between theoretical predictions and empirical patterns may stem from subsequent evolutionary changes in life history traits and responses to local environmental conditions.

### Bridging theory and empirical results

Two scenarios could potentially reconcile our theoretical predictions with observed data. First, one might hypothesize that sampled diploid populations are predominantly outcrossing while polyploid populations are predominantly selfing, an arrangement that would naturally produce lower evolvability in polyploids. However, this scenario contradicts previous studies on *E, incanum*, which rather show that polyploidy is associated with a shift toward more outcrossing phenotypes. For example, changes in *Erysimum incanum* in flower size and spatial position of sexual organs related to genome duplication tend to be more fitted to outcrossing strategies (García-Muñoz et al., 2023).

A more plausible explanation is that polyploidy alters the selective landscape (the ω^2^ parameter in our simulations), by modifying the ecology of populations. Polyploidy is associated with periods of environmental turmoil and contemporary stressful conditions (see Van de Peer et al. 2021 for a review), and harsher conditions are known to decrease the evolvability of populations (Pujol and Pannell, 2008; Martinez-Padilla et al., 2017; Pennington et al., 2021). Our data support this interpretation: while mountain environments generally enhanced evolvability, likely due to increased ecological heterogeneity, the interaction between polyploidy and environments was negative, especially for hexaploids. These populations, located in more arid southern regions, appear to experience harsher environmental constraints. Polyploid lineages can undergo rapid niche differentiation (Baniaga et al., 2020; Padilla-García et al., 2023), and adaptation to narrow, extreme environments may reduce within-population genetic variance. In contrast, diploid and tetraploid populations inhabit more variable habitats, potentially retaining broader evolutionary potential. A summary of the study’s results and the current literature is available in Table 6.

**Table 6.**
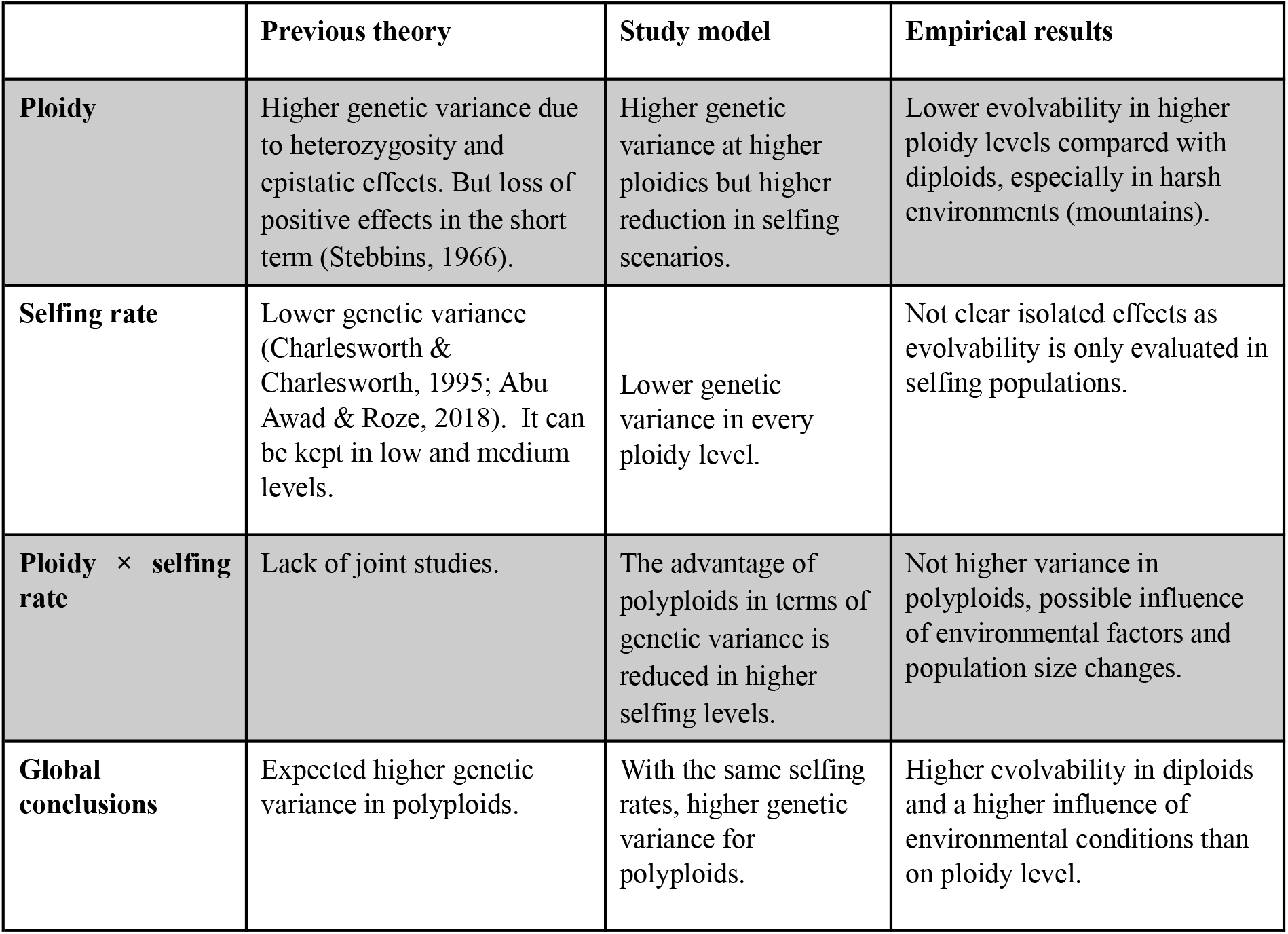
Comparative summary of effects of ploidy and mating system in genetic variance according to previous knowledge, the here developed mathematical model, and empirical results of our study.

|  | **Previous theory** | **Study model** | **Empirical results** |
| --- | --- | --- | --- |
| **Ploidy** | Higher genetic variance due to heterozygosity and epistatic effects. But loss of positive effects in the short term (Stebbins, 1966). | Higher genetic variance at higher ploidies but higher reduction in selfing scenarios. | Lower evolvability in higher ploidy levels compared with diploids, especially in harsh environments (mountains). |
| **Selfing rate** | Lower genetic variance (Charlesworth & Charlesworth, 1995; Abu Awad & Roze, 2018). It can be kept in low and medium levels. | Lower genetic variance in every ploidy level. | Not clear isolated effects as evolvability is only evaluated in selfing populations. |
| **Ploidy × selfing rate** | Lack of joint studies. | The advantage of polyploids in terms of genetic variance is reduced in higher selfing levels. | Not higher variance in polyploids, possible influence of environmental factors and population size changes. |
| **Global conclusions** | Expected higher genetic variance in polyploids. | With the same selfing rates, higher genetic variance for polyploids. | Higher evolvability in diploids and a higher influence of environmental conditions than on ploidy level. |

### Transgressive segregation and polyploidy

Our results also challenge the assumption that polyploidy promotes transgressive segregation. Polyploidy levels were negatively associated with transgressive segregation, while mountain conditions had a positive effect (as shown by TS2 and the variance ratio; Table 3). In contrast, TS1 revealed no significant effects. This suggests that polyploidy does not necessarily increase the likelihood of generating novel phenotypes beyond the parental range, despite theoretical predictions of enhanced genetic covariance in polyploids (Clo, 2022b), and empirical evidence of previous studies suggesting that increased heterozygosity in polyploids leads to novel allele combinations and diversity (Sattler et al., 2015; De los Reyes, 2019).

Interestingly, common garden conditions were associated with reduced transgressive segregation, while greenhouse conditions appeared to enhance it. This highlights the role of environmental complexity in shaping the emergence of novel trait combinations. Although polyploids often display broader phenotypic ranges (Čertner et al., 2019), their adaptive potential may still be greater in more variable environments, despite the observed reduction in transgressive segregation under certain conditions.

### Conclusion and perspectives

In summary, our empirical findings suggest that diploids are more evolvable than polyploids and that ploidy level has limited influence on transgressive segregation. These results contrast with theoretical expectations that posit increased genetic variance with higher ploidy. One likely explanation lies in differences in selective pressures, as hexaploids tend to live in more arid and extreme environments and have adapted to more restricted conditions than their diploid and tetraploid counterparts.

Our current model does not incorporate non-additive genetic effects, such as dominance or epistatic interactions. While dominance reduces the difference in adaptive potential among mating strategies in diploids, its role in polyploids has not yet been explored. Neutral models suggest that the contribution of dominance variance in polyploids should be minor (Gallais, 2003) and only transient (Walsh, 2005). So, including dominance should not have changed our predictions qualitatively. Epistasis could have a more pronounced effect depending on whether ploidy increases or not the number of epistatic interactions compared to diploids, and on the directionality of epistasis (Carter et al., 2005). No trivial conclusions can be drawn, suggesting a direction for future investigation.

Refining the model to distinguish between neo- and established polyploids, and to incorporate events such as bottlenecks, would also provide a better understanding of the adaptive potential of polyploids and selfing populations.

Future models should distinguish between neo- and established polyploids and include demographic processes such as founder effects or bottlenecks to better reflect real-world scenarios. Although theoretical models provide valuable general frameworks, they cannot fully account for the complexities of natural populations. Empirical data are therefore essential to refine and improve these models.

Our results reinforce the importance of combining theoretical and empirical approaches to better understand how mating systems and ploidy jointly influence the adaptive potential of plant populations. This integration is crucial for predicting evolutionary trajectories under environmental change, especially in diverse and dynamic systems such as *Erysimum incanum*.

## Acknowledgements

We thank editors and reviewers for comments that improved the clarity of the manuscript. D.D. is supported by a Ph.D. studentship from the Hauts-de-France region. J.C. is supported by the CNRS. Experimental work was supported by a grant from the Organismo Autónomo de Parques Nacionales *globalHybrids* [Ref: 2415/2017], and the Ministerio de Ciencia e Innovación projects *OUTevolution* [PID2019-111294GB-I00/SRA/10.13039/501100011033] and *Meenerva* [PID2022-139405OB-I00/AEI /10.13039/501100011033], including FEDER funds. AJM-P was funded partially by the European Commission under the Marie Sklodowska-Curie Action Cofund 2016 EU agreement 754446 and the UGR Research and Knowledge Transfer—Athenea3i. COC was supported by the *Contratos Predoctorales (FPU) Universidad de Granada-Banco Santander* in the 2021 call. AG-M was supported by the *OUTevolution* project [PID2019-111294GB-I00/SRA/10.13039/501100011033].

## Competing interests

We have no competing interests to declare.

## Author contributions

A.G.M and C.F. performed the experimental work and data collection. M.A. and A.J.M.P. did the experimental design and got the funding for its development. C.O.C. performed the statistical analyses with the support of J.C. D.D. performed the theoretical analyses with the support of J.C. C.O.C., D.D., M.A., and J.C. wrote the original draft, all authors commented and approved the final version of the manuscript.

## Data availability

The simulation model is available at: https://github.com/DianeDouet/ploidy_mating_QG. The experimental dataset generated during the current study is available from the corresponding author on reasonable request.

## Notes

https://github.com/DianeDouet/ploidy_mating_QG

### Competing Interest Statement

The authors have declared no competing interest.

### Summary of Updates

The text has been modified to add clarifications and explanations based on the reviews.

