## Supplementary material for "The effects of ploidy and mating system on the evolvability of populations: theoretical and empirical investigations"

**Table S1. Sample sizes for individuals in families across different environmental, ploidy, and heterozygosity levels were obtained in the experiment.**

| Selfing | Diploid | Tetraploid | Hexaploid |
|---|---|---|---|
| GREENHOUSE | 16 in 9 | 172 in 35 | 130 in 24 |
| COMMON GARDEN | 150 in 16 | 80 in 13 | 102 in 12 |
| MOUNTAIN | 91 in 23 | 205 in 45 | 20 in 10 |
| Outcrossing | | | |
| GREENHOUSE | 4 in 2 | 21 in 6 | 49 in 8 |
| COMMON GARDEN | 49 in 4 | 48 in 6 | 63 in 8 |

**Table S2.** Evolvabilities of individual and flower traits in *Erysimum incanum* for each ploidy and environment level (GH: Greenhouse; CG: Common garden; MT: Mountain).

| | Diploid | | | Tetraploid | | | Hexaploid | | |
|---|---|---|---|---|---|---|---|---|---|
| | GH | CG | MT | GH | CG | MT | GH | CG | MT |
| **Plant height** | 0.714 | 1.483 | 1.108 | 3.335 | 0.179 | 3.646 | 1.594 | 1.293 | 0.610 |
| **Stalk diameter** | 0.995 | 3.551 | 5.349 | 7.561 | 2.996 | 14.089 | 6.736 | 3.068 | 1.578 |
| **Number of flowers** | 0.198 | 1.126 | 1.633 | 5.130 | 1.083 | 4.654 | 2.803 | 0.488 | 1.314 |
| **Petal length** | 1.477 | 3.145 | 3.049 | 4.351 | 1.251 | 2.671 | 3.120 | 0.682 | 0.219 |
| **Corolla diameter** | 0.781 | 3.490 | 8.453 | 2.894 | 1.113 | 4.428 | 1.087 | 0.508 | 0.294 |
| **Corolla tube length** | 1.697 | 2.463 | 1.797 | 3.462 | 1.391 | 1.315 | 1.995 | 2.669 | 0.256 |
| **Long filament length** | 0.862 | 1.902 | 2.229 | 1.591 | 0.260 | 0.727 | 0.328 | 0.246 | 0.276 |
| **Short filament length** | 1.601 | 4.230 | 3.315 | 2.253 | 0.311 | 1.699 | 0.770 | 0.528 | 0.186 |
| **Style length** | 1.728 | 1.986 | 385.974 | 2.317 | 0.409 | 1.211 | 0.711 | 0.967 | 0.424 |
| **Herkogamy** | 0.574 | 0.803 | 364.565 | 1.426 | 0.496 | 1.195 | 1.899 | 2.578 | 0.932 |

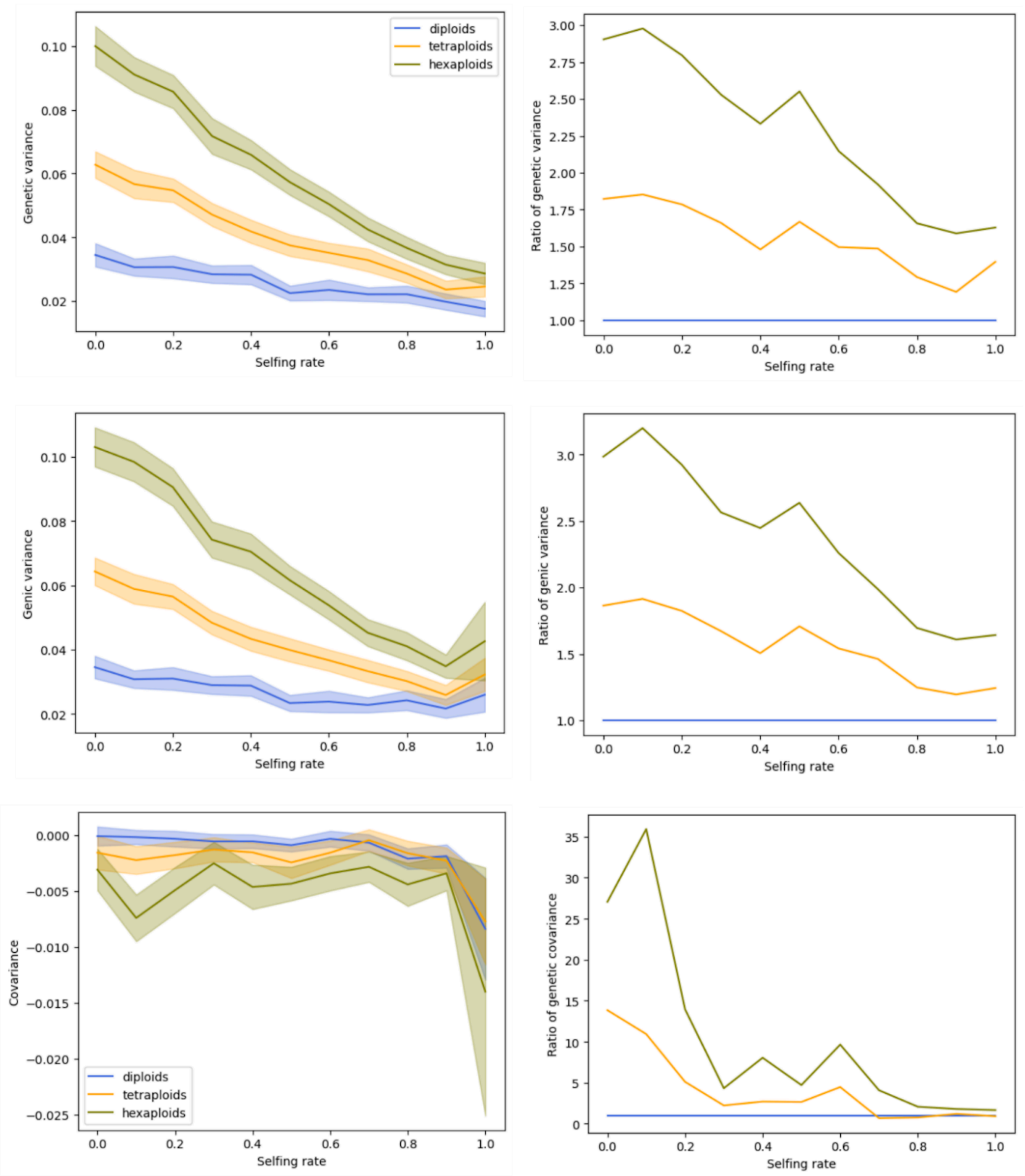


**Figure S1.** Genetic variance, genic variance and covariance (left) and their ratios between polyploid and diploid populations (right) as functions of the selfing rate for different levels of ploidy, when $U = 0.005$ and $\omega^2 = 1$.

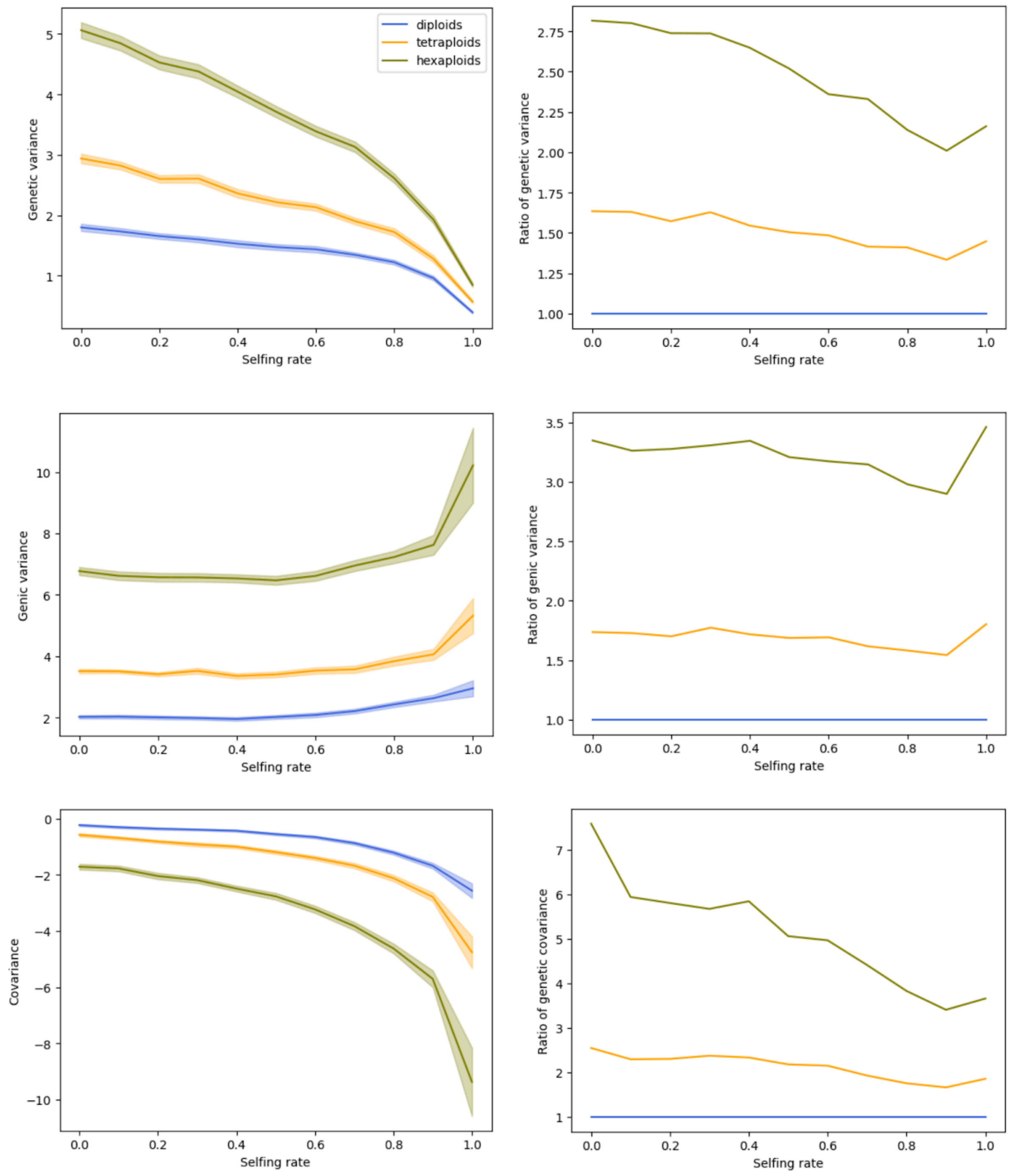


**Figure S2.** Genetic variance, genic variance and covariance (left) and their ratios between polyploid and diploid populations (right) as functions of the selfing rate for different levels of ploidy, when $U = 0.1$ and $\omega^2 = 9$.

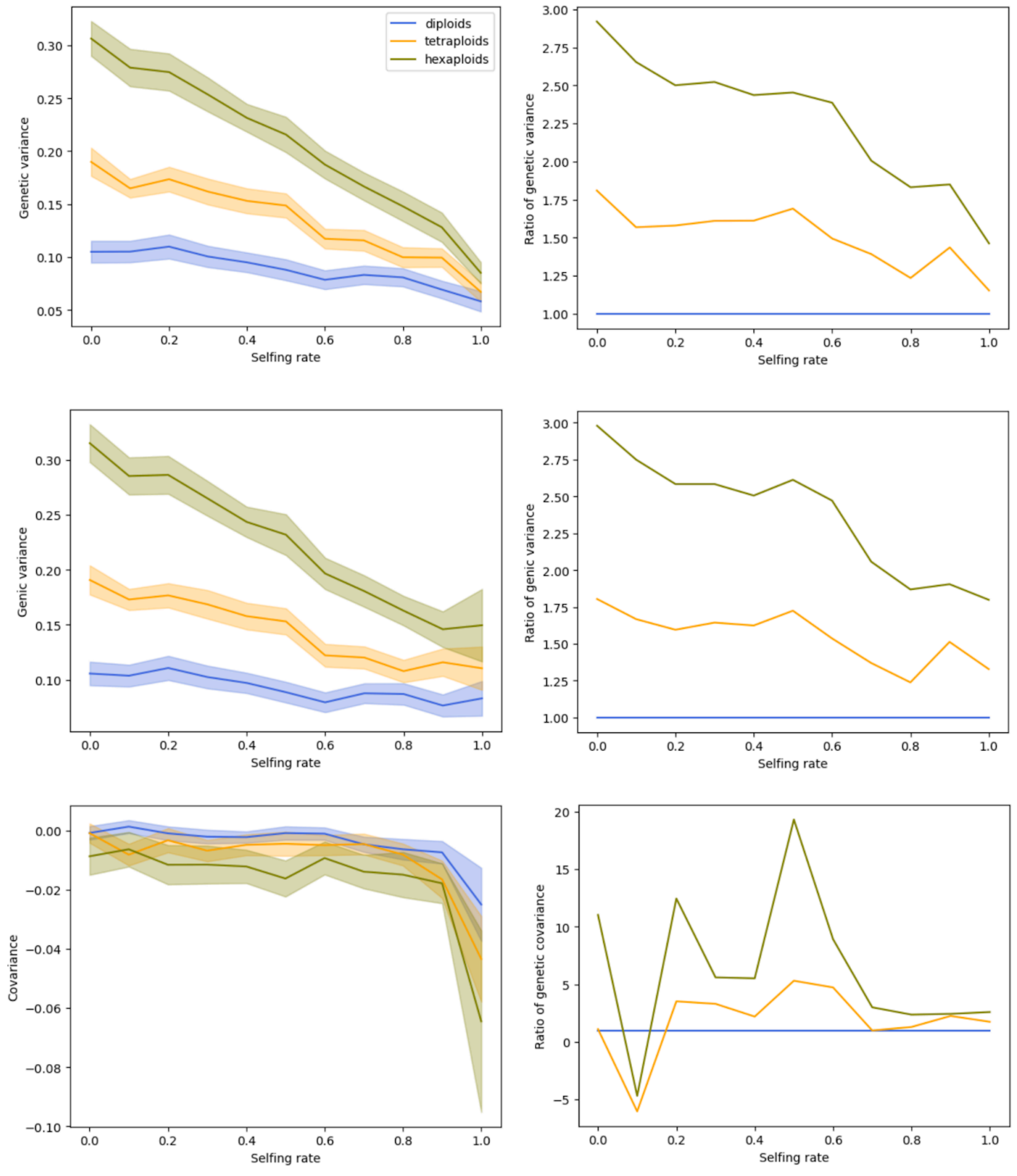


**Figure S3.** Genetic variance, genic variance and covariance (left) and their ratios between polyploid and diploid populations (right) as functions of the selfing rate for different levels of ploidy, when $U = 0.005$ and $\omega^2 = 9$.
